# Biomolecular condensates of Chlorocatechol 1,2-Dioxygenase as enzymatic microreactors for the degradation of polycyclic aromatic hydrocarbons

**DOI:** 10.1101/2023.05.29.542454

**Authors:** Nathan N. Evangelista, Mariana C. Micheletto, Luis F. S. Mendes, Antonio J. Costa-Filho

**Author notes:** Corresponding author: Antonio J. Costa-Filho, Av. Bandeirantes 3900, FFCLRP/USP, Monte Alegre, Ribeirão Preto, SP, Brazil, 14040-901.

## Abstract

Polycyclic aromatic hydrocarbons (PAHs) are molecules with two or more fused aromatic rings that occur naturally in the environment due to incomplete combustion of organic substances. However, the increased demand for fossil fuels in recent years has increased anthropogenic activity, contributing to the environmental concentration of PAHs. The enzyme chlorocatechol 1,2-dioxygenase from *Pseudomonas putida* (Pp 1,2-CCD) is responsible for the breakdown of the aromatic ring of catechol, making it an interesting player in bioremediation strategies. Pp 1,2-CCD can tolerate a broader range of substrates, including halogenated compounds, than other dioxygenases. Here, we report the construction of a chimera protein able to form biomolecular condensates with potential application in bioremediation. The chimera protein was built by conjugating Pp 1,2-CCD to low complex domains (LCDs) derived from the DEAD-box protein Dhh1. We showed that the chimera could undergo liquid-liquid phase separation (LLPS), forming a protein-rich liquid droplet under different conditions (variable protein and PEG8000 concentrations and pH values), in which the protein maintained its structure and main biophysical properties. The condensates were active against 4-chlorocatechol, showing that the chimera droplets preserved the enzymatic activity of the native protein. The possible application of these microreactors in bioremediation is discussed.

## INTRODUCTION

Polycyclic aromatic hydrocarbons (PAHs) are molecules containing two or more bonded aromatic rings. They naturally occur in the environment due to the incomplete combustion of organic substances, e.g., forest fires and volcanic activity. The increasing demand for fossil fuels in the last decades has escalated the contribution of anthropogenic activity to the environmental PAH concentration [1,2]. Moreover, world oil markets have rebounded from the massive demand shock after the Covid pandemic [2,3]. In this context, more than a hundred known PAHs have been identified. The Environmental Protection Agency of the United States (US EPA) has cited sixteen as priority contaminants because of their oncogenic, teratogenic, and mutagenic characteristics [4]. These are detrimental to human health and the environment. They are recalcitrant and resistant to degradation due to their stable and consistent nature.

Many technologies are available for the remediation of polluted sites. Bioremediation and catalytic reaction are green techniques that can degrade the toxic pollutants in the environment, converting them into non-toxic or less harmful compounds [5]. A bacterial genus of *Pseudomonas* can degrade PAHs of different molecular masses, from simpler aromatic compounds, such as naphthalene, to compounds such as benzo(a)pyrene (both present in the US EPA list) [6,7]. The bacteria’s ability for bioremediation is related to enzymes that transform the PAH complex molecules into common intermediate compounds of their metabolic routes to be used as extensive carbon and energy sources [8].

The metabolic funnel of PAH degradation begins with the oxidation of one of the benzene rings by an oxygenase enzyme via the incorporation of atmospheric oxygen [8]. The oxidation of the first benzene ring is usually the rate-limiting step in the degradation of PAHs [8]. Subsequent oxidations lead to the formation of the common intermediate, catechol, or its halogenated derivatives. The product of the intradiol cleavage (between carbons 1 and 2) of catechol is *cis,cis*-muconic acid, which can be converted to adipic acid when hydrogenated. Adipic acid is an important industrial compound used, for example, in producing benzene-fnylon-6,6 [9,,11].

The chlorocatechol 1,2-dioxygenase enzyme from *Pseudomonas putida* (Pp 1,2-CCD) is responsible for the fission of the aromatic ring of catechol. It plays a crucial role in the processes of bioremediation and sustainable production of adipic acid. Pp 1,2-CCD has tolerance to a greater diversity of substrates than other dioxygenases, such as catechol 1,2-dioxygenase and protocatechuate 3,4-dioxygenase [12]. In addition to the catechol, Pp 1,2-CCD can cleave halogenated substrates such as 3-chlorocatechol, 4-chlorocatechol, 4-fluorcatechol, 3,5-dichlorocatechol, 3,5-dibromocatechol, tetrachlorocatechol, protocatechuate, 3-methylcatechol, 4-methylcatechol, 3-methoxycatechol, and 4-nitrocatechol [13].

The Pp 1,2-CCD catalytic site has a Fe^3+^ ion coordinated by four residues: Tyr130, Tyr164, His188, and His190 in a trigonal bipyramidal geometry. Akin to other members of the same family, Pp 1,2-CCD functional unit is homodimeric [9]. A hydrophobic tunnel crosses the structure from one face to the other between the two dimer subunits, where a phospholipid molecule, possibly a phosphatidylinositol (PI), is attached [9,13,14]. Previous results from our group demonstrated that the presence of this molecule exerts an allosteric control of the enzymatic activity, decreasing its affinity for the substrate and increasing the initial maximum reaction rate [14]. Regarding the enzymatic kinetics, our group has also demonstrated that Pp 1,2-CCD does not behave according to conventional Michaelis-Menten kinetics [10,13], with the product binding at the enzyme’s catalytic site. Inhibition by the product should be related to cell homeostasis mechanisms to avoid the toxicity associated with high concentrations of *cis,cis*-muconic acid [10,14].

In this context, the wide substrate diversity of Pp 1,2-CCD makes it an interesting target for applications in bioremediation. Using environmental conditions to control the Pp 1,2-CCD activity would be an adequate strategy for applying the enzyme in bioremediation. Immobilization techniques have become convenient tools for modifying enzymes’ biochemical/biophysical properties, such as activity, stability, specificity, selectivity, and decreased inhibition [15, 16]. The protein immobilization also allows enzyme reuse and facilitates the separation of catalyst and product [15]. However, enzyme immobilization has been a challenge in biotechnology. Although allowing greater control of the reaction, many immobilization techniques can cause denaturation or changes in the structure due to the immobilization procedures, leading to the loss of enzymatic activity [17]. On the other hand, increasing the durability of the enzyme and preventing inhibition by the products should be tackled in the search for an efficient application [15].

One alternative to better control the enzyme’s reaction conditions and applicability in biotechnology is Liquid-Liquid Phase Separation (LLPS). LLPS is a thermodynamically driven, reversible phenomenon that demixes two liquids into two distinct liquid phases with different solute concentrations [18,19,20]. The equilibrium between mixing and demixing strongly depends on many physicochemical parameters, including solute concentration, temperature, pressure, pH, and crowding agents [21,22,23].

A known strategy to induce LLPS of otherwise globular enzymes consists of attaching Low Complexity Domains (LCDs) to the termini of the target enzyme [24]. LCDs have ubiquitous flexibility and structural features without a unique or defined structure due to their amino acid sequence, playing central roles in protein recognition, interactions, and phase separation [18,25]. These domains are poor in complexity with repeated or low-diverse amino acid sequences [25,26]. LCDs have the property to induce protein LLPS under certain conditions, making them versatile tools with applications in several fields, such as food engineering [27], enzyme technology [28], and water treatment [29].

The conjugation of LCDs to a given protein forms a chimera protein, which, due to the presence of the LCDs, is capable of undergoing LLPS. In this study, a novel Pp 1,2-CCD chimera protein, conjugated with LCDs based on the DEAD-box protein Dhh1, was constructed and characterized. The potential use of these microreactors for bioremediation applications is discussed.

## RESULTS

### Construction and characterization of the Pp 1,2-CCD chimera

The Pp 1,2-CCD enzyme (hereafter also named CCDWT) had been previously characterized in our group [10,14,30] and served as the starting point of our chimera construction. We built a chimera protein (CCD-LCD_2_) by conjugating Pp 1,2-CCD to molecular adhesives based on the DEAD-box protein Dhh1 sequence [31]. The LCDs used here were those described in FALTOVA et al. [24] and have been reported to induce phase separation when used as molecular adhesives in globular proteins [24,31]. In order to obtain this chimera enzyme, we built a modified pET28a vector containing restriction sites for BamHI and HindIII conjugated to the coding gene of the LCDs. The gene for Pp 1,2-CCD was isolated using Polymerase Chain Reactions (PCR) and inserted in the restriction sites for BamHI and HindIII using a subcloning process. The expression and purification of the chimera enzyme followed the protocols described by ARAUJO et al. [32] and resulted in high-yield and pure samples. The success of these processes was confirmed by SDS-PAGE and Size Exclusion Chromatography (SEC) (Figure S1). The SDS-PAGE of the expressed product showed pure proteins and the expected molecular masses for both chimera (44 kDa) and wild-type (28 kDa) proteins.

We then analyzed potential conformational changes in the CCD core of the chimera protein that could affect the enzyme activity. The amino acid sequences of CCDWT and CCD-LCD_2_ were submitted to AlphaFold2 analyses. AlphaFold2 is an artificial intelligence-based software designed to predict protein structure from the protein’s amino acid sequences [33]. The results are shown in Figure 1A, where it is possible to notice similar structures of the globular domain of the CCDWT and the chimera. The two conjugated intrinsically disordered domains are also seen in the latter. Moreover, the Root-Mean Square Deviation (RMSD) of the superimposed globular structures of CCD-LCD_2_ and CCDWT was 0.118 Å, which suggests low differences between the two proteins. These results indicate that conjugation did not alter the enzyme structure. Therefore, activity should be preserved.

**Figure 1.**
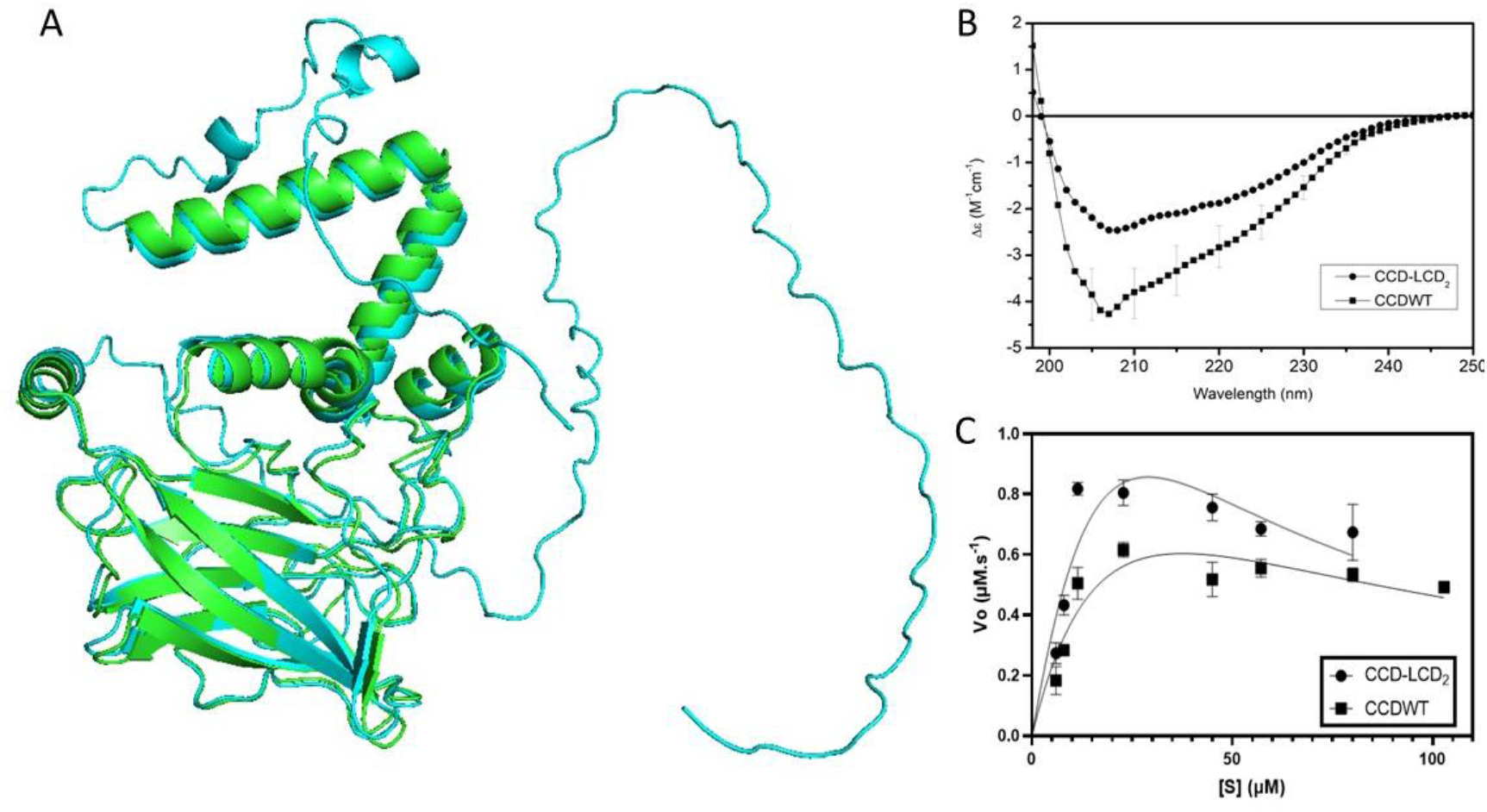
Construction and characterization of the CCD-LCD_2_. (A) 3D structures for CCDWT (green) and CCD-LCD_2_ (blue) predicted using Alphafold2. (B) CD spectra for CCDWT (squares) and CCD-LCD_2_ (circles). (C) Enzymatic activities for CCDWT (squares) and CCD-LCD_2_ (circles).

Following the structural analysis using AlphaFold2, we performed circular dichroism (CD) experiments to probe the protein’s secondary structure experimentally. CD data also suggested that the proteins share a similar fold despite the LCD additions in the chimera protein (Figure 1B). Both CD spectra present a similar profile consisting of a negative peak around 208 nm and a shoulder around 222 nm, as expected for proteins containing α-helices [34]. However, the chimera enzyme showed an overall decrease in the peak intensities. This decrease and the poorer resolution of the peaks in the CD spectrum of the chimera could be due to the mixture of contributions to the final CD spectrum coming from the disordered LCDs present in the chimera structure. When analyzing the insertion of the LCDs using AlphaFold2 (Figure 1A), it is possible to notice a significant *tail*-like structure at the N-and C-terminals. As CD spectra are based, in principle, on the secondary structure contents, our results likely reflect differences in optical absorption of circularly polarized light arising from the combination of a globular domain (CCD) with the intrinsically disordered regions of the LCDs.

The next step in characterizing the CCD-LCD_2_ chimera was to assess its enzymatic activity, as described in Materials and Methods. Figure 1C shows that CCWWT and CCD-LCD_2_ presented similar enzymatic activities. The chimera protein could reach a higher V_max_ than its wild-type version (3.78 μM.s^-1^ for CCD-LCD_2_ and 1.44 μM.s^-1^ for CCDWT). We previously showed that chlorocatechol 1,2-dioxygenase is an enzyme that does not follow Michaelis-Menten (MM) kinetics [10]. Instead, it does show a product inhibition mechanism that can be described by adjusting the regular MM scheme to include an inhibition term in the velocity equation [10,14]. Thus, using this adapted product inhibition mechanism, we inferred the affinity of the substrate for the active site in both cases (CCDWT and CCD-LCD_2_) based on the calculation of the K_m_ values. We observed that the affinity thus determined was higher for CCDWT, with K_m_ equal to 26.0 μM, compared to CCD-LCD_2,_ which showed a K_m_ of 49.4 μM. The inhibition constant (K_i_) was 17.0 μM for the chimera and 54.4 μM for the wild-type version. The values of K_m_ and K_i_ indicate that the chimera enzyme had a lower affinity for the substrate and was less affected by the formation of the reaction product.

### Chimera proteins can undergo Liquid-Liquid Phase Separation (LLPS) under certain conditions

After the initial biophysical characterization, we studied the LLPS propensity of the chimera protein CCD-LCD_2_ by Differential Interference Contrast Microscopy (DIC) and Dynamic Light Scattering (DLS). The increase in macromolecular crowding has been shown to play a critical role in inducing the liquid-liquid phase separation of many proteins [35]. Therefore, we used the crowding agent PEG8000 to test the LLPS propensity of the chimera protein. CCD-LCD_2_ showed liquid droplet formation at pH 8.0 and in the range of 2% up to 10% m/v of the crowding agent (Figure 2A). At first, a very high protein concentration (12 g/L) in the standard working buffer (20 mM Tris-HCl, 250 mM NaCl) was used to guarantee enough intermolecular interactions to support the phase separation. These initial results were used to guide the subsequent experiments that established in which general conditions CCD-LCD_2_ presented phase separation.

**Figure 2.**
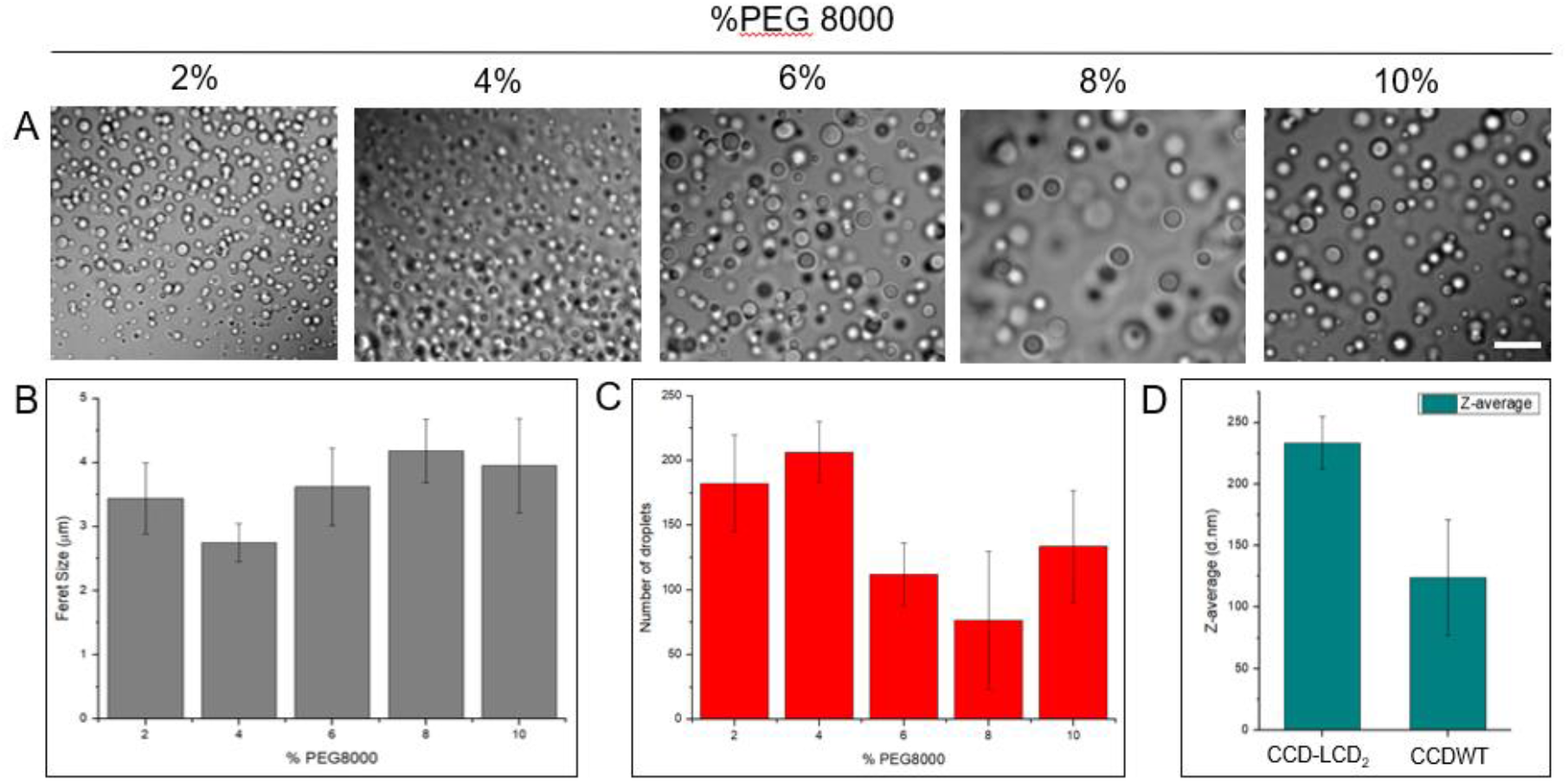
(A) DIC images showing the droplets formed by CCD-LCD2 as a function of PEG concentration. Scale bar: 10 μm. (B) Estimated Feret size for each PEG concentration. (C) Number of droplets per area of 4.10^3^ μm². (D) Hydrodynamic radius estimated by DLS for the chimera and wild-type proteins at 10% PEG 8000. All the measurements were performed in a 20 mM Tris-HCl buffer at pH 8.0 and in the presence of 250 mM NaCl. The protein concentration was 12 g.L^-1^.

The images obtained by DIC microscopy in Figure 2A were treated using ImageJ software [36], and the number of droplets per image and the Feret size were estimated. Figure 2B and C show that the Feret did not significantly change, whereas the number of droplets decreased at 6% PEG8000 concentration and remained constant. These trends are likely related to the stabilization of the condensates at high concentrations of PEG8000. In less crowded environments, the molecules are more dispersed, and the water-protein interactions are favored instead of the protein-protein contacts. However, at higher concentrations of PEG8000, the protein is forced to have more protein-protein contacts, thus phase separating. The scenario remains until there are insufficient protein-protein contacts in the dilute phase to form more condensates. Based on the data shown in Figure 2, it is possible to state that the chimera protein can undergo phase separation from low to high concentrations of PEG8000, and the images are better collected and easily visualized in more crowded environments. Therefore, the chosen standard condition for phase separation of CCD-LCD_2_ was 10% PEG8000.

Having determined an optimized condition regarding the crowding agent, we further investigated CCD-LCD_2_’s capability of undergoing LLPS as a function of pH changes. Since pH is a critical parameter for protein stabilization and its variations may induce protein denaturation, we first monitored how pH affected the secondary structures of CCD-LCD_2_ and CCDWT. Figure 3 shows the CD spectra measured for the proteins as a function of pH, at 10% PEG concentration, and with the protein in conditions expected for the dispersed phase (low protein concentration, as in Figure 4). The results showed that, upon decreasing the pH value to 6.0, the CD spectra of the proteins lost intensity, likely due to a combined effect of sample precipitation (visible in the cuvette) and protein denaturation. CD spectral deconvolution using BEstSel [37] allowed us to quantify the secondary structure content in each case. Table 1 shows that, at pH 6.0, the turns and other secondary elements responded for ca. 75% of the total secondary structure content, with no contribution from α-helices. There were still β-sheets at pH 6.0, but that should be analyzed with caution since the CD spectra of β-structures are variable and could thus be used by the deconvolution algorithm to fit the remaining negative signal at pH 6.0.

**Table 1.** Secondary structure content of CCDLCD_2_ and CCDWT at pH 6.0, 7.0, and 8.0 estimated using BestSel software [34].

| *Secondary structure* | <u>CCDWT (%)</u> |  |  | <u>CCD-LCD<sub>2</sub> (%)</u> |  |  |
| --- | --- | --- | --- | --- | --- | --- |
|  | pH 8.0 | pH 7.0 | pH 6.0 | pH 8.0 | pH 7.0 | pH 6.0 |
| *α-Helix* | 20.9 | 13.8 | 1.3 | 16.2 | 12.1 | 0 |
| *Antiparallel* | 14.9 | 25.7 | 23.3 | 18.8 | 27.1 | 27.2 |
| *Parallel* | 4.6 | 3.8 | 0 | 4.6 | 1.6 | 0 |
| *Turn* | 8.5 | 12.1 | 19.7 | 11.8 | 12.1 | 19.6 |
| *others* | 51.2 | 44.6 | 55.7 | 48.6 | 47 | 53.3 |

**Figure 3.**
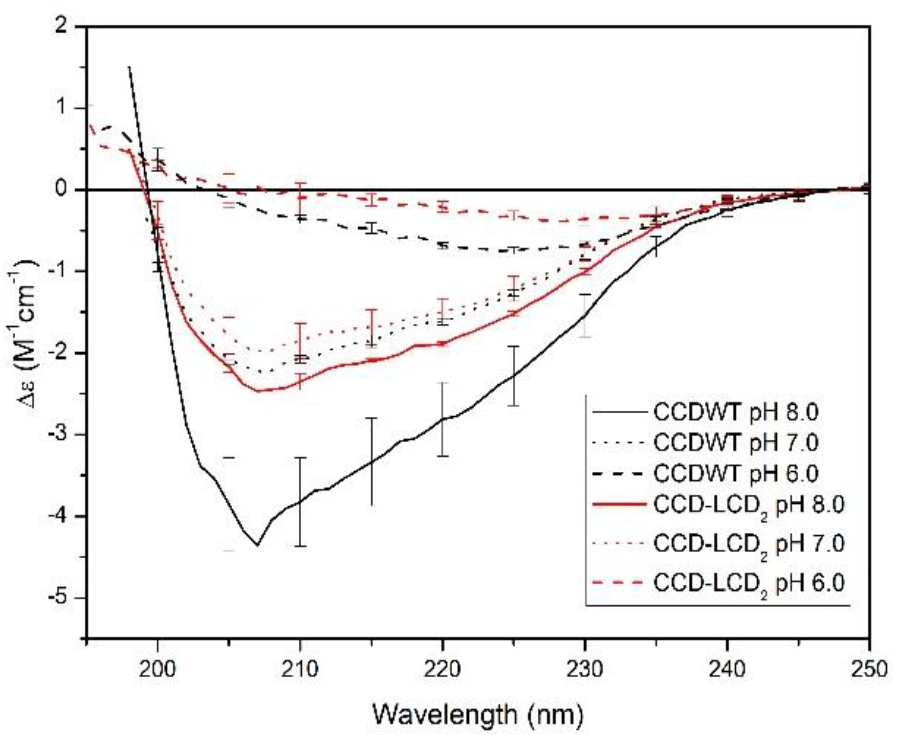
Effects of pH on the secondary structure of the proteins dispersed in a 10% PEG8000 solution. The CD spectra exhibit a protein secondary structure loss as the pH decreases. At pH 6.0, wild-type and chimera have very weak signals, indicating possible denaturation.

**Figure 4.**
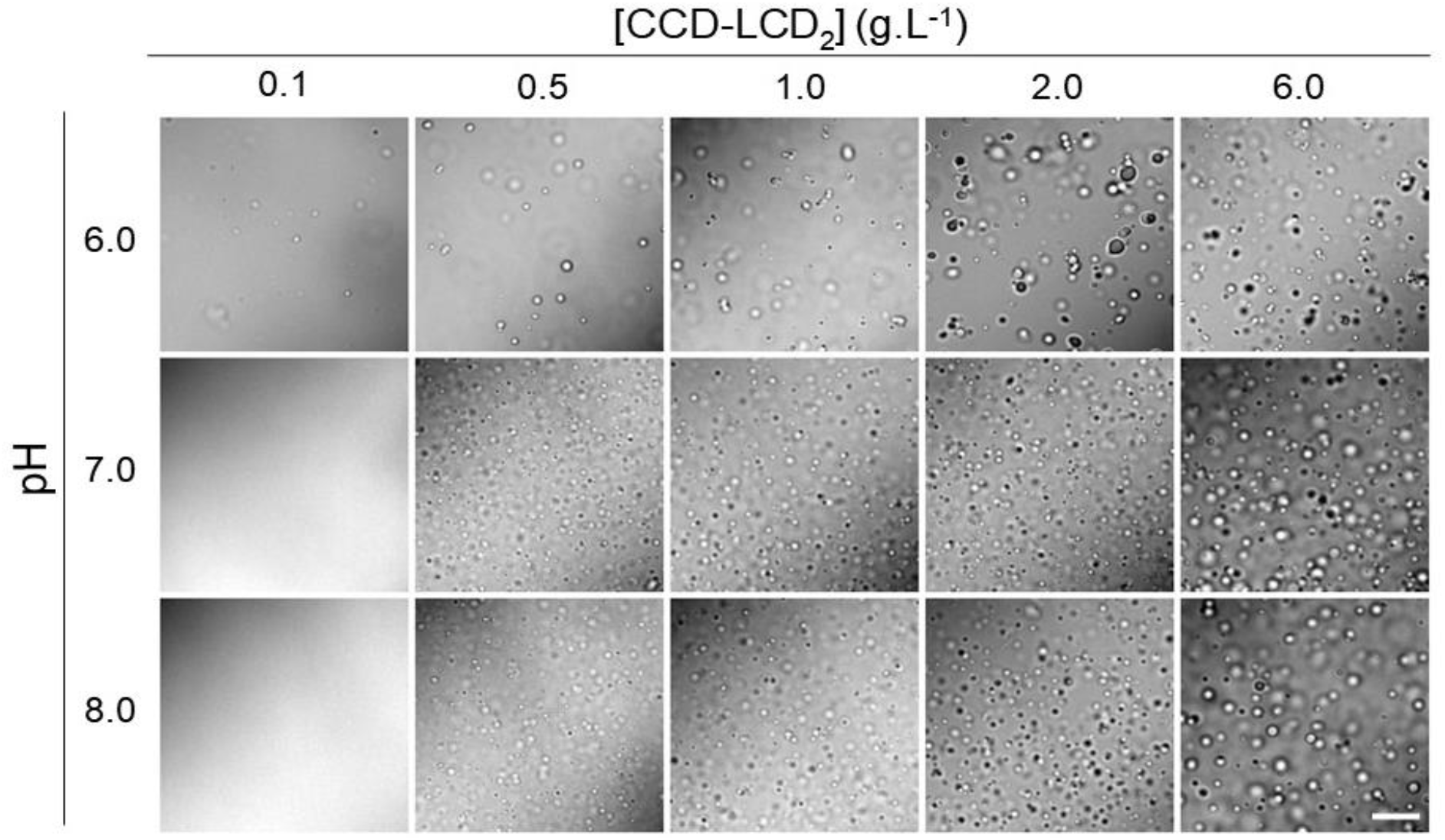
Phase diagram built using DIC images of CCD-LCD_2_. All images were obtained in a 50 mM phosphate buffer and 250 mM NaCl, with 10% PEG 8000 addition. Scale bar: 10 μm.

After analyzing the pH effect on the protein’s structure, a phase diagram was constructed, varying the pH value and protein concentration to determine the conditions for which phase separation occurred. Figure 4 shows the DIC images obtained for CCD-LCD_2_ at pH 6.0, 7.0, and 8.0 and upon varying the protein concentration. We can see that, at pH 7.0 and 8.0, droplets were readily formed in high amounts even at the lower protein concentrations (0.5 g/L, which corresponds to ca. 10 mM). There was a threshold in protein concentration to observe droplet formation. The droplets were mobile condensates, and fusion events were observed at pH 7.0 and 8.0. At pH 6.0, although droplets could still be seen, sample precipitation occurred, in agreement with our CD experiments. Furthermore, droplets presented more irregular non-spherical shapes with lower mobility in solution and no fusion events were found, indicating a potential solid-like state.

### CCD-LCD_2_ droplets are porous and can evolve to a mature and irreversible state with solid-like properties

The maturation and reversibility of the CCD-LCD_2_ droplets were also studied to gain information on their main rheological properties. Maturation is related to the capability of the protein to phase separate and evolve to gel- and solid-like condensates, which present different rheological properties, such as the absence of fusion events (due to high viscosity properties, for example) or extinction of the dynamic equilibrium between the protein in the dilute and condensed phases.

The reversibility of the phase separation was initially tested by varying protein concentration upon different incubation times. We observed that the liquid-liquid phase separation induced by PEG8000 (as described before) could be reversed by decreasing protein concentration to values lower than 0.5 g/L (Figure 4). Figure 5A shows that droplets formed in the presence of 10% PEG8000 (incubation time of 20 minutes) could be dissolved upon diluting the protein to 0.1 g/L after another 20-minute interval. The liquid-like properties of the droplets could also be perceived in the fusion events shown in Figure 5C. Here, samples incubated at shorter times could coalesce, as seen in images with a 30-second screenshot interval. On the other hand, droplets incubated at room temperature for 24 hours evolved to a state with no fusion events, loss of mobility, and unable to dissolve upon protein dilution (0.1 g.L^-1^) (Figure 5B). These clearly indicate that the droplet evolved to a more solid-like condensate.

**Figure 5.**
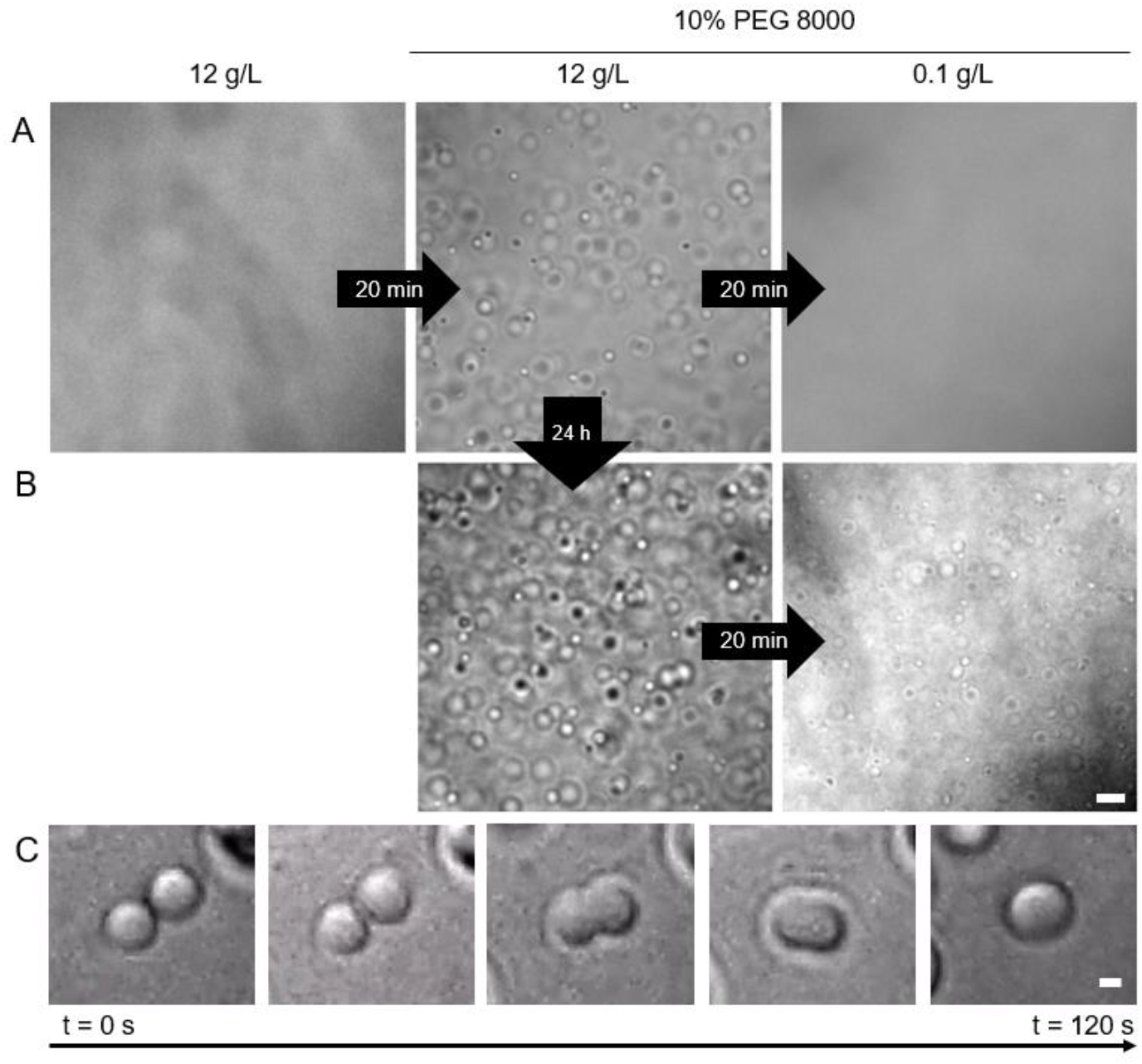
Droplet reversibility and coalescence. Maturation of CCD-LCD_2_ condensates followed at different incubation times. The phase separation was induced by 10% PEG8000 in 20 mM Tris-HCl buffer, 250 mM NaCl, pH 8.0, and at high protein concentration. (A) When incubated for short periods, the droplets showed liquid-like properties and were highly dynamic. These structures were reversible and could evolve to a soluble phase after dilution. (b) After 24 hours, irreversible condensates were formed. Scale bar: 5 μm. (C) Coalescence event seen in images taken at a 30 s time interval. Scale bar: 1 μm.

Besides the droplet reversibility and coalescence, we also investigated their permeability using an assay that consisted in adding the Sypro Orange fluorescent probe to the droplet solution and monitoring their permeability with time. Sypro orange has a high affinity for hydrophobic sites and, when bound to those sites, experiences an increase in its quantum yield, reflected in an increase in its fluorescence. Therefore, we used Sypro orange as a probe to check possible flow into the droplets. Sypro Orange was added to a 5 μL CCD-LCD_2_ droplet solution, and images were taken after incubation. To observe the incorporation of the probe, we monitored the fluorescence intensity increasing inside the droplets using Steady-state Fluorescence Microscopy. The results after the addition are presented in Figure 6 and clearly show the permeability of the probe into the droplets after 40 minutes.

**Figure 6.**
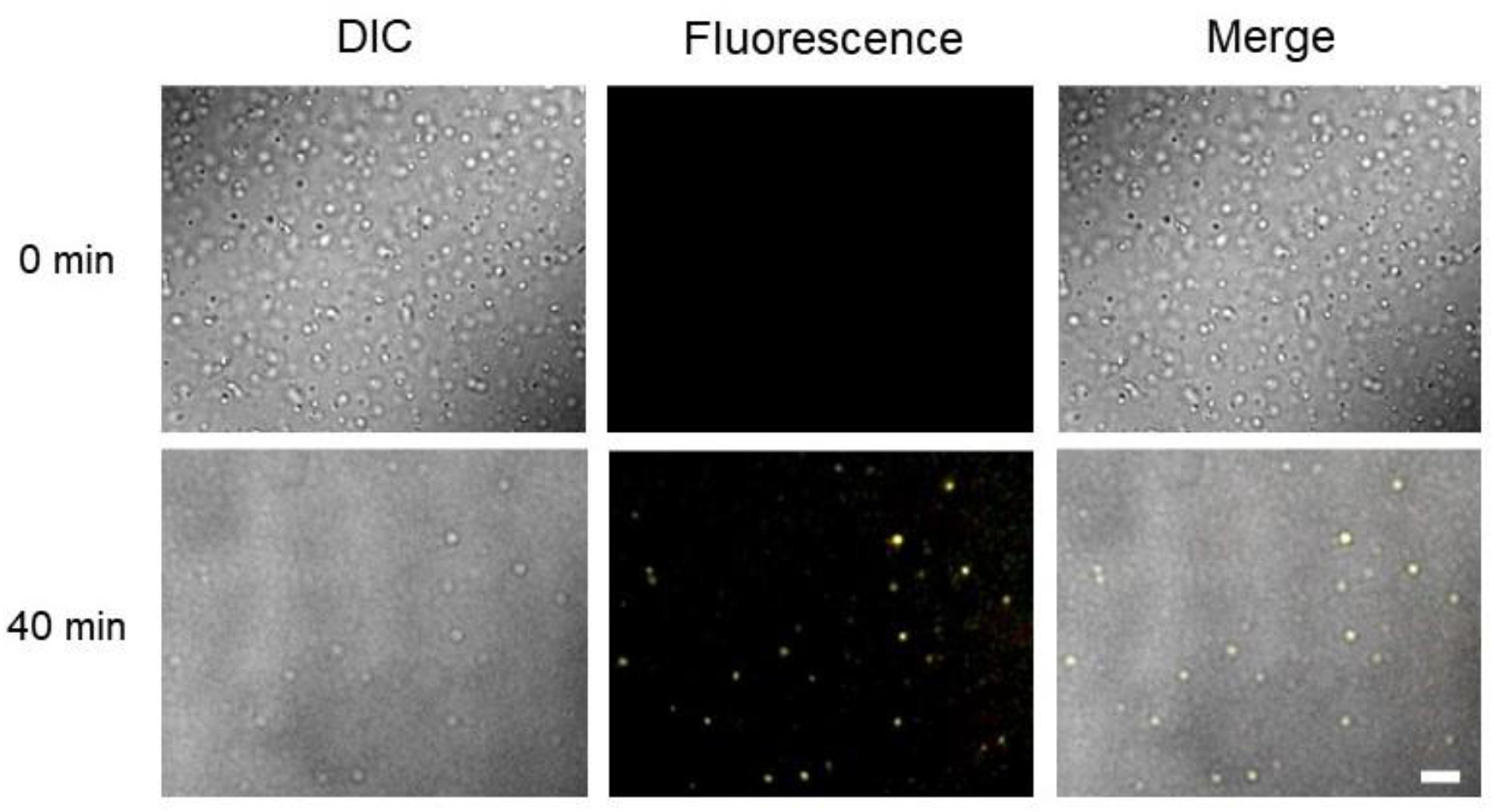
CCD-LCD_2_ condensates are permeable to Sypro orange, which could diffuse to the droplet interior, as shown by the increased fluorescence within the droplet after 40 minutes. DIC images were used to locate the droplets, and fluorescence to locate Sypro orange within the droplets. Merged images are in the right-most panels. All images were measured in a 20 mM Tris-HCl buffer, at pH 8.0 and in the presence of 250 mM NaCl and 10% PEG8000. Scale bar: 5 μm.

### CCD-LCD_2_ droplets maintain the enzyme’s native activity

Lastly, we tested the droplets as enzymatic microreactors, able to perform reactions in a membrane-less compartmentalized space. The enzymatic activity assays performed with CCD-LCD_2_ not in LLPS conditions showed that the chimera enzyme kept the native activity of CCDWT, indicating that the addition of the LCDs did not modify the overall capacity of the enzyme to catalyze the reaction (Figure 1).

To perform the assays with CCD-LCD_2_ in the condensates, the phase separation was initially induced with 25 μM CCD-LCD_2_ in Tris-HCl buffer at pH 8.0 in the presence of 250 mM NaCl and 10% PEG8000. Here, we tested the conversion of 4-chlorocatechol to *cis,cis*-muconic acid in phase separation conditions, where most protein molecules are not free in solution but rather in the protein-rich droplet phase. We used substrate concentrations ranging from 25 μM up to 100 μM. The catalysis was monitored by measuring the absorbance spectrum of a sample containing 4-chlorocatechol before and after the addition of CCD-LCD_2_. The reactions were performed at room temperature for 10 seconds, and the concentration of the product was estimated by its molar extinction coefficient, as described in the Materials and Methods section. The results are presented in Figure 7 and show that the initial absorbance spectrum of 4-chlorocatechol, with its typical maximum around 286 nm, vanished upon the addition of CCD-LCD_2_ along with the appearance of a strong absorbance around 260 nm, which is characteristic of *cis-cis*-muconic acid. The assay, thus, showed the total conversion of the substrate 4-chlorocatechol to the product *cis,cis*-muconic acid, confirming the catalytic activity of CCD-LCD_2_ in LLPS conditions.

**Figure 7.**
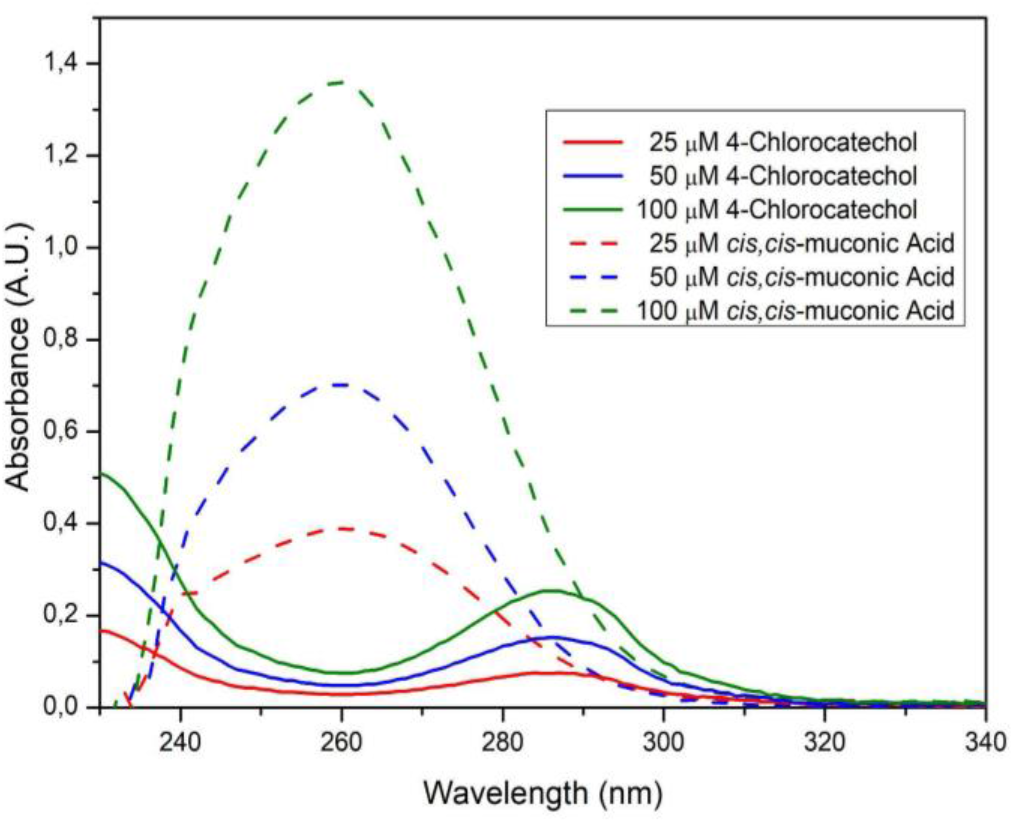
Absorption spectra for different concentrations of 4-chlorocatechol and its degradation product. Solid lines indicate samples containing no enzyme but 4-chlorocatechol. Dash lines indicate the product formed by the reaction in the presence of chimera CCD-LCD_2_, after 10 seconds of incubation. All the substrate is suggested to react at short periods, even at high concentrations. The enzymatic assays were performed in Tri-HCl buffer at pH 8.0, in the presence of 250 mM NaCl and PEG 8000 addition. The presence of liquid protein droplets was confirmed by microscopy before and after the reaction.

## DISCUSSION

Our results report the construction of a chimera protein able to phase separate and successfully catalyze its original reaction. The conditions triggering LLPS were determined, with the possibility of phase separation at >0.5 g.L^-1^ protein concentrations and at neutral or low alkaline solutions. Polluted environments suitable for treatments via bioremediation strategies involving enzymes from the catechol dioxygenase family usually present neutral acid conditions, as well are related to the presence of a variety of organic salts, heavy metal ions (commonly oxidized by combustion) and, as expected, polyaromatic hydrocarbons (PAHs) [38,39]. The presence of organic salts and compounds creates aggressive and crowded conditions for proteins due to the high concentration of those molecules as particulate matter (PM) [38]. In our work, we proposed using a chimera protein that can catalyze reactions even upon those aggressive conditions and act on the degradation of high PAH concentrations (above 14,45 mg.L^-1^ or 1,445% w/v) at very short times (less than 10 seconds). According to Malaswka and Wiołkomirski (2001) [39], we can consider soil as heavily polluted at ranges above 1% w/w of pollutant concentration. Therefore, we can state that we could produce microreactors able to degrade highly polluted soils, as the improvement of a protein able to catalyze these reactions at restricted sites and, finally, with the maintenance of its original structure even at aggressive conditions.

## CONCLUSIONS

The possibility of using biomolecular microreactors is a great advantage to perform and control reactions inside a specific site. Here, we have developed a chimera protein able to undergo phase separation in the presence of PEG 8000 to a protein-rich phase, maintaining its globular domain structure and main biophysical properties. The total secondary structure content was conserved, and the substrate affinity was maintained. We showed that the generated droplets are highly dynamic and have liquid-like properties, as shown by their coalescence and permeability. In addition, these droplets can also be functionalized as microreactors and act on the degradation of pollutants. In future perspectives, we want to demonstrate that the droplets can make enzymatic cascades to increase the efficiency of PAHs metabolization.

## MATERIALS AND METHODS

### DNA construction, protein expression and purification

A pET28a modified vector was constructed using the LCD sequence (derived from Dhh1 from *Saccharomyces cerevisiae*, AA 1-47 and AA 426-508). This sequence was synthesized and optimized for expression in E. coli by Exxtend. The LCD vector was fused with CCD through a subcloning process. The recombinant proteins were produced using LB medium and E. coli BL21 (DE3) competent cells (Novagen). The bacterial culture was grown to 0.8 O.D. at 37ºC under 200 rpm agitation. Then the cells are induced to expression using 0.5 mM of isopropyl-β-D-thiogalactoside (Sigma Aldrich) and incubated for 16 hours at 18ºC and 200 rpm agitation. The recombinant proteins were purified using nickel affinity chromatography (Qiagen). As the final purification step, we performed size exclusion chromatography using a Superdex 10/300 GL on an Akta Purifier System (GE Healthcare). We used Tris-HCl buffer pH 8.0 with a strength of 20 mM and added 250 mM NaCl. Protein concentration was estimated using ProtParam software, with an extinction coefficient of 34,505 M^-1^.cm^-1^ and 45,965 M^-1^.cm^-1^ for CCDWT and CCD-LCD_2_, respectively. The success of each step has been checked by SDS-PAGE electrophoresis in a Bio-Rad Mini-PROTEAN® Tetra Vertical Electrophoresis Cell. Proteins were stored at 4ºC.

### Circular Dichroism (CD)

Far UV-CD spectra were measured using a Jasco J-810 spectrometer (Jasco Corporation, Japan) using a quartz cell with 0.1 cm path length. The spectra were recorded from 250 to 190 nm, with a scanning speed of 100 nm/min and a protein concentration of 0.1 g.L^-1^.

### Liquid-liquid phase separation (LLPS) induction and droplets analysis

The addition of PEG 8000 induced the LLPS. Each solution was incubated for 20 minutes, and images were taken using an IX71 inverted microscope (Olympus). The buffers used for the samples were 20 mM Tris-HCl pH 8.0, 20 mM phosphate buffer pH 7.0, and 20 mM phosphate buffer pH 6.0, each with 250 mM NaCl addition. A U-MWB2 mirror unit with excitation BP460/490 was used for steady-state fluorescence microscopy studies. 5 μL of the solution was placed onto a glass slide and taken to the microscope at 20ºC. All the images were analyzed through ImageJ software [36].

### Permeability studies

In order to visualize Sypro Orange incorporation by protein droplets, 5 μL protein solution at 2 g.L^-1^ was placed onto a glass slide. The microscope camera was positioned, and 1 μL Sypro Orange was placed into a different position. After 40 minutes, images were taken to verify the incorporation of the fluorescent probe by the droplets.

### Enzymatic activity assays

Enzymatic activity was measured using an Ultrospec 2100 pro spectrophotometer using a quartz cell with 1 cm pathlength. We monitored the absorbance in 260 nm (the maximum absorption for the product *cis,cis*-muconic acid) for 200 seconds and determined the initial reaction rate for different substrate concentrations. Activity analysis and curve fitting were made by Graphpad Prism 6, using the inhibition model known for the wild-type enzyme. To monitor product formation at LLPS conditions, absorption spectra were initially taken for a solution containing no enzyme and 4-chlorocatechol at different concentrations. The reaction was performed for 10 seconds, and the spectra of each solution were again measured. All analyses were made on the software Graphpad Prism6 following the substrate inhibition model as previously described by MELO [10].

### Dynamic Light Scattering

The hydrodynamic radius was measured using a Zeta Sizer instrument (Malvern, UK) for both proteins, at a concentration of 12 g.L^-1^ in a 20 mM Tris-HCl pH 8.0 buffer, with 250 mM NaCl addition.

## ACKNOWLEDGEMENTS

The authors acknowledge Fundação de Amparo à Pesquisa do Estado de São Paulo (FAPESP) for the financial support (Grant nos. 2015/50366-7 and 2020/15542-7). NNE, MCM, and LFSM thank FAPESP for their fellowships (Grant nos. 2020/01152-2, 2018/13016-6, and 2017/24669-8). AJCF thanks Conselho Nacional de Desenvolvimento Científico e Tecnlógico (CNPq) for the partial financial support (Grant no. 306682/2018-4).

## Notes

### Competing Interest Statement

The authors have declared no competing interest.

