## Supplementary figure for "Biomolecular condensates of Chlorocatechol 1,2-Dioxygenase as enzymatic microreactors for the degradation of polycyclic aromatic hydrocarbons"

1- Laboratório de Biofísica Molecular, Departamento de Física, Faculdade de Filosofia, Ciências e Letras de Ribeirão Preto, Universidade de São Paulo, Ribeirão Preto, SP, Brazil.

2- Grupo de Biofísica Molecular Sérgio Mascarenhas, Departamento de Física e Ciência Interdisciplinar, Instituto de Física de São Carlos, Universidade de São Paulo, São Carlos, SP, Brazil.

Figure S1: Results of the protein expression and purification protocols: (A) SDS-PAGE. (B) Size exclusion chromatography.

A

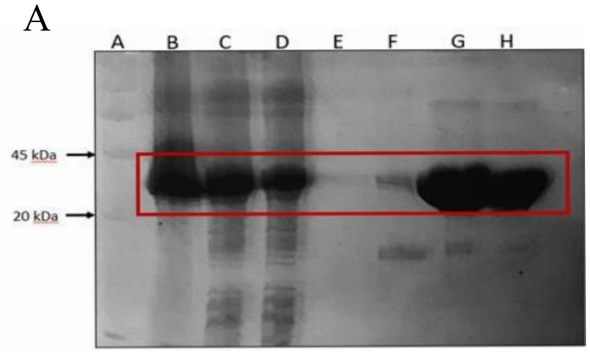


B

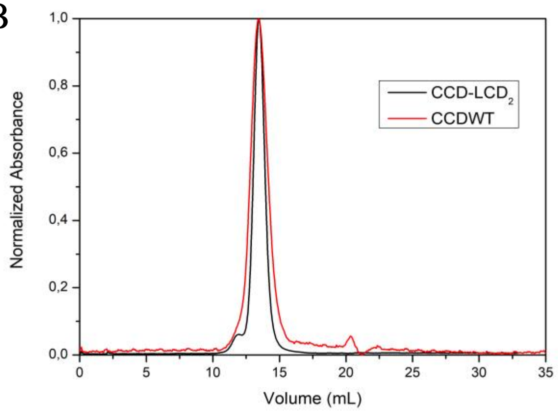
